# Ketamine disrupts gaze patterns during face viewing in the common marmoset

**DOI:** 10.1101/2021.02.16.431438

**Authors:** Janahan Selvanayagam, Kevin D. Johnston, Raymond K. Wong, David J. Schaeffer, Stefan Everling

## Abstract

Faces are stimuli of critical importance for primates. The common marmoset (*Callithrix jacchus*) is a promising model for investigations of face processing, as this species possesses oculomotor and face processing networks resembling those of macaques and humans. Face processing is often disrupted in neuropsychiatric conditions such as schizophrenia (SZ) and thus it is important to recapitulate underlying circuitry dysfunction preclinically. The N-Methyl-D-aspartate (NMDA) non-competitive antagonist ketamine has been used extensively to model the cognitive symptoms of SZ. Here, we investigated the effects of a subanesthetic dose of ketamine on oculomotor behaviour in marmosets during face viewing. Four marmosets received systemic ketamine or saline injections while viewing phase-scrambled or intact videos of conspecifics’ faces. To evaluate effects of ketamine on scan paths during face viewing, we identified regions of interest in each face video, and classified locations of saccade onsets and landing positions within these areas. A preference for the snout over eye regions was observed following ketamine administration. In addition, regions in which saccades landed could be significantly predicted by saccade onset region in the saline but not the ketamine condition. No significant drug effects were observed for phase-scrambled videos. Effects on saccade control were limited to a reduction in saccade amplitudes during viewing of scrambled videos. Thus, ketamine induced a significant disruption of scan paths during viewing of conspecific faces but limited effects on saccade motor control. These findings support the use of ketamine in marmosets for investigating changes in neural circuits underlying social cognition in neuropsychiatric disorders.

## Introduction

The common marmoset (*Callithrix jacchus*), is a rapidly emerging non-human primate model for neuroscientific research. This species possesses a lissencephalic cortex which is advantageous for fMRI (Hung et al., 2015a; Schaeffer et al., 2020), laminar electrophysiology (Johnston et al., 2019) and optical imaging (Kondo et al., 2018; Sadakane et al., 2015). Marmosets have also convergently evolved a rich social behavioural repertoire that mirrors that of humans, including pair bonding and alloparental care (Schiel & Souto, 2017). Sophisticated multi-modality social communication has additionally been observed in this species and includes a library of distinct vocalizations, scent marking, tactile communication such as social grooming, and visual communication via body postures, physical gestures, and facial expressions (Bezerra & Souto, 2008; de Boer et al., 2013; Kemp & Kaplan, 2013; Moreira et al., 2013). Together, this combination of practical experimental advantages and natural social behaviours is ideal for studies of the neural basis of social cognition, and impairments in processing of social signals often observed in neuropsychiatric conditions (Dodell-Feder et al., 2015; Feuerriegel et al., 2015; Yager & Ehmann, 2006)

Facial processing is of critical importance to social cognition in primates. The ability to detect, identify and extract social information from faces is highly efficient and supported by a specialized network of cortical and subcortical areas exhibiting selectivity for faces (Freiwald, 2020; Johnson, 2005). Investigations in humans have shown selective activation in the fusiform gyrus, lateral occipital cortex, superior temporal sulcus and inferotemporal cortex (Haxby et al., 2000). Similarly, in macaques, distinct “face patches” can be observed along the occipito-temporal axis (Tsao et al., 2008). Electrophysiological investigations of these areas reveal neurons that are not only face-selective, but also sensitive to specific features of individual faces such as view direction and facial identity (Chang & Tsao, 2017; see for review Freiwald, 2020; Freiwald et al., 2009; Freiwald & Tsao, 2010). In the marmoset, recent fMRI evidence has revealed a face network comparable to that observed in macaques and humans (Hung et al., 2015a, 2015b; Schaeffer et al., 2020), and previous behavioural investigations have shown that marmosets use gaze information and facial expressions for social communication in both head-free and head-restrained contexts in the laboratory (Kemp & Kaplan, 2013; Mitchell et al., 2014). Such cross-species similarities in face processing and face networks suggest that the marmoset has considerable potential as a model species in face processing research.

Irregularities in face processing are a common characteristic of some human neuropsychiatric conditions. Patients with schizophrenia, for example, have been shown to exhibit impairments in configural face processing (Baudouin et al., 2008; Joshua & Rossell, 2009; Shin et al., 2008) aberrant facial emotion processing (see for review Morris et al., 2009), and abnormal gaze patterns during face viewing, including avoidance of the eyes (Loughland et al., 2002; Phillips & David, 1998; Williams et al., 1999). Neural correlates of these impairments have been documented, including attenuated amplitudes for event related potential components associated with face processing (Feuerriegel et al., 2015; McCleery et al., 2015), and structural and functional abnormalities in face processing areas such as the fusiform gyrus (P. J. Johnston et al., 2005; Onitsuka et al., 2006; Quintana et al., 2003). and emotional information processing areas such as the amygdala (Hall et al., 2008; Holt et al., 2006; see for review Aleman & Kahn, 2005).

The N-Methyl-D-aspartate (NMDA) non-competitive antagonist ketamine has long been used to model symptoms of schizophrenia (Krystal, 1994). Unlike dopaminergic agents, which primarily model the positive symptoms of schizophrenia (e.g., hallucinations, delusions), the ketamine model additionally produces negative symptoms including flat affect, emotional withdrawal, and cognitive symptoms (see for review Beck et al., 2020). These cognitive symptoms include impairments of configural and emotional face processing (Lundin et al., 2020; Neill et al., 2015; Reed et al., 2019; Schmidt et al., 2013), suggesting that investigations of face processing using the ketamine model may provide a gateway to further understanding of cognitive and face-processing impairments in schizophrenia.

Investigations in non-human primate models have provided valuable insight into the neural basis of face processing, as tools like intracortical microstimulation and pharmacological manipulations have been used to disrupt the system and investigate causal relationships (Afraz et al., 2006; Moeller et al., 2017; Sadagopan et al., 2017). For example, pharmacological manipulations have demonstrated a causal role of the posterior superior temporal sulcus in gaze following behaviour, which relies on configural face processing, in the macaque (Roy et al., 2014). Here, we evaluated the utility of the common marmoset as a model for cognitive impairments affecting face processing by monitoring eye movements during a simple face-viewing task following administration of ketamine or saline. Ketamine administration altered the pattern of saccades between facial features with minimal impacts on oculomotor behaviour in general. Taken together, our findings show that investigations of oculomotor behaviour in marmosets can provide valuable insights into cognitive functions such as face processing and how these may be impacted in disease states.

## Methods

### Subjects

Data were collected from 4 adult common marmosets (*Callithrix jacchus*; 1 female; weight 382– 615 g; age 29 – 74 months). All experimental procedures conducted were in accordance with the Canadian Council of Animal Care policy on the care and use of laboratory animals and a protocol approved by the Animal Care Committee of the University of Western Ontario Council on Animal Care. The animals were under the close supervision of university veterinarians.

Prior to these experiments, all 4 marmosets underwent an aseptic surgical procedure to implant a combination recording chamber/head restraint, the purpose of which was to stabilize the head during eye tracking experiments. The chamber implantation procedure is described in detail in Johnston et al. (2018). Subsequently, each animal was acclimated to restraint in a custom designed primate chair (Johnston et al., 2018; Schaeffer et al., 2019).

### Data collection

Marmosets were seated in a custom primate chair (Johnston et al., 2018) with the head restrained, inside a sound attenuating chamber (Crist Instruments Co., Hagerstown, MD, USA). A spout was placed at the animals’ mouth to deliver reward (acacia gum) via an infusion pump (Model NE-510, New Era Pump Systems, Inc., Farmingdale, New York, USA). Eye position was calibrated in each session by rewarding 300 to 600 ms fixations on dots presented centrally or at +/- 5° abscissa or ordinate on the display monitor using the CORTEX real-time operating system (NIMH, Bethesda, MD, USA). All stimuli were presented on a CRT monitor (ViewSonic Optiquest Q115, 76 Hz non-interlaced, 1600 x 1280 resolution). Eye positions were digitally recorded at 1 kHz via video tracking of the left pupil (EyeLink 1000, SR Research, Ottawa, ON, Canada). Animals were intermittently rewarded at random time intervals to maintain their interest.

#### Injections

Following calibration, each marmoset received an intramuscular injection of either 0.5ml/kg of saline or ketamine (dose=1mg/kg). Each marmoset completed two sessions on separate days, with the order of saline and ketamine injections counterbalanced. Data collection commenced 5 minutes following injection.

#### Stimuli

The same block design used in (Schaeffer et al., 2020) was used here, in which monkeys viewed short video clips consisting of conspecifics’ faces or scrambled version of these faces (12s), interleaved with fixation blocks (18s) where they were presented with a central, circular, black fixation stimulus (2°, see Figure 1). Marmosets were not required to fixate during these blocks. During video blocks, the fixation stimulus was removed, and the video was presented at the center of the screen (16.5° height x 29.5° width) with face videos and their scrambled versions being presented in a pseudorandomized manner, counterbalanced across subjects. For the face videos, 12s clips were created from videos of 4 marmosets seated in a marmoset chair (iMovie, Apple Incorporated, California, USA). Scrambled versions of the videos were created by random rotation of the phase information, preserving motion components by using the same random rotation matrix for each frame (Matlab, The Mathworks, Matick, MA). Stimuli were presented via Keynote (Version 7.1.3, Apple Incorporated, California, USA) with stimulus timing achieved using a TTL pulse emitted by a photodiode. Animals were intermittently rewarded at random time intervals to maintain their interest.

**Fig 1.**
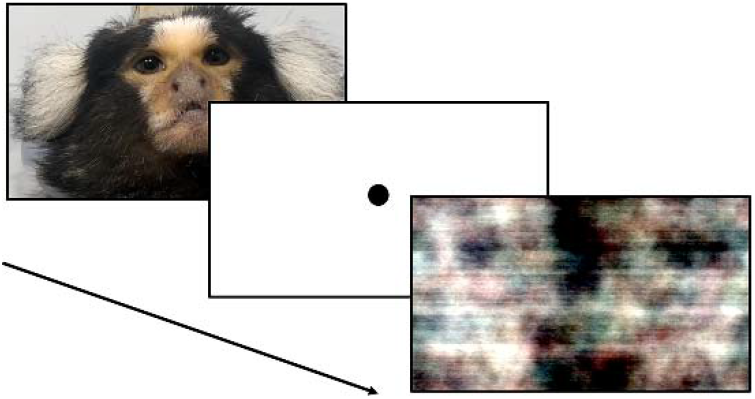
Task Design. Face Video and Scrambled Video blocks (12s) were presented in a pseudorandomized order, interleaved with Fixation blocks (18s).

### Data analysis

Analysis was performed using Python code written in-house. Eye velocity (visual deg/s) was obtained by smoothing and numerical differentiation. Saccades were defined as radial eye velocity exceeding 30 deg/s for a movement greater than 0.5 visual degrees. Fixations were defined as periods where radial eye velocity remained below 10 deg/s for at least 50 ms. Rectangular regions of interest (ROI) were manually labelled in the video clips at a subset of ‘key’ frames and linearly interpolated using Python code such that the ROI outlined facial features of interest (e.g., eyes, snout) as they moved in the video clip. Accuracy of the interpolation was then manually verified, adding additional key frames as necessary.

Differences in saccade amplitudes and fixation durations between conditions were evaluated using 2 x 3 repeated measures analyses of variance (ANOVA) with factors Drug Treatment (saline, ketamine) and Viewing Block (Fixation, Scrambled Video, Face Video). These were carried out in SPSS (v.25, IBM Corp, 2019). Greenhouse-Geisser corrections were applied where the assumptions of sphericity were violated. Partial eta squared (*η_p_^2^*) is reported as a measure of effect size. Post-hoc tests of means were corrected using the Bonferroni method. Differences in transition probabilities between regions of interest between treatments were analyzed with multinomial logistic regressions using the multinom function (nnet package v7.3-14, Ripley, 2020) and custom code written in R (R v3.5.2, R Core Team, 2013).

## Results

### Saccade amplitude

Median saccade amplitudes were computed for each monkey (Figure 2A). A repeated measures ANOVA was conducted for the Viewing Blocks (Fixation, Scrambled Video, Face Video) and drug Treatments (Saline, Ketamine) on these data (see Figure 2a). We observed no significant main effect of Viewing Block, *F*(2,6) = 2.83,*p* = .136, *η_p_^2^* = .486, a significant effect of Treatment, *F*(1,3) = 10.96,*p* = .045, *η_p_^2^* = .785, and a significant Viewing Block x Treatment interaction, *F*(2,6) = 12.74,*p* = .007, *η_p_^2^* = .809, showing that the effect of Treatment was significant only in the Scrambled Video Block (Δ = 2.54 degrees, *p* = .021) but not in the Fixation (Δ = 1.85 degrees, *p* = .064) or Face Video (Δ = 1.30 degrees, *p* = .099) Blocks.

**Fig 2.**
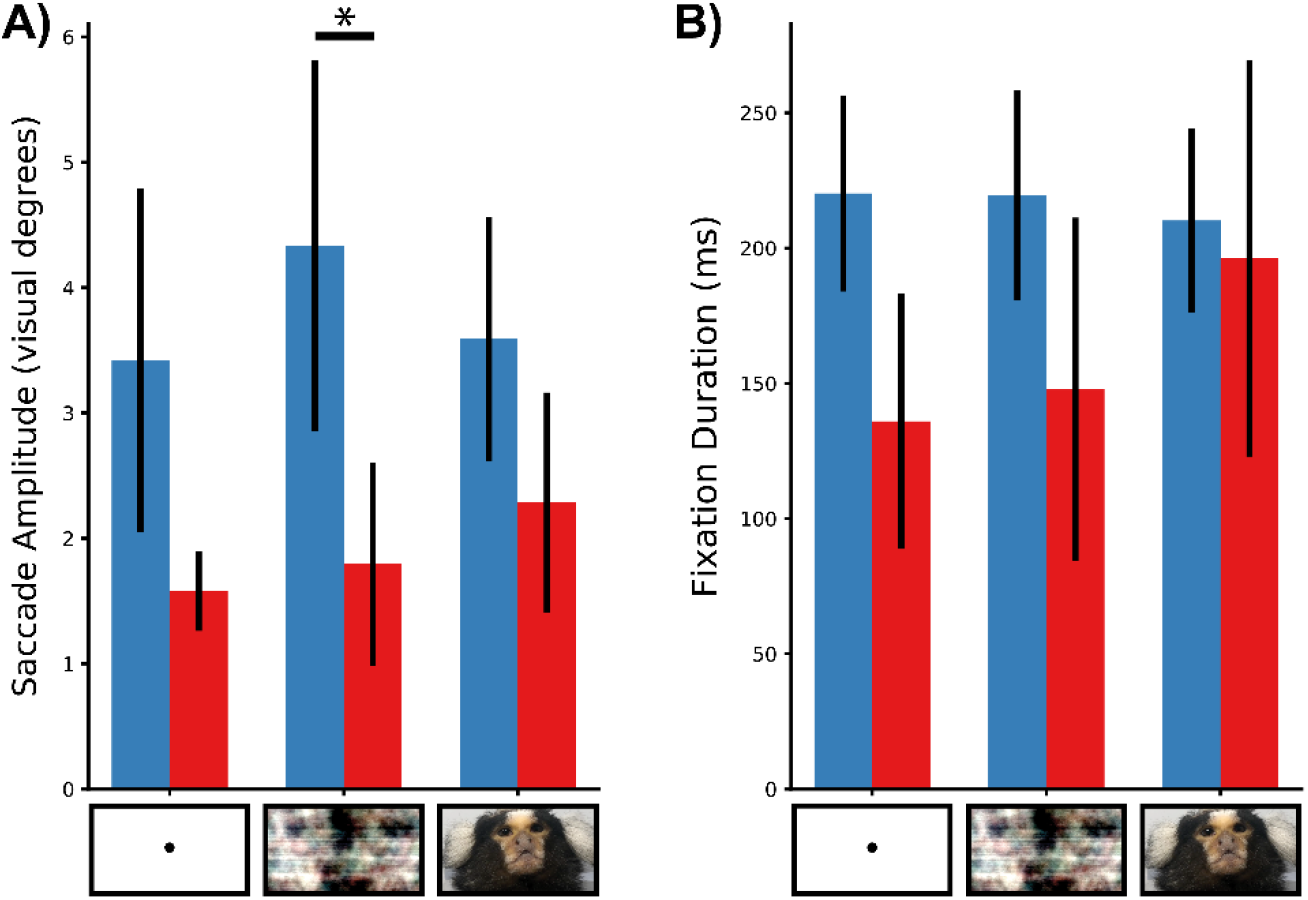
Ketamine effects on saccade amplitudes and fixation durations during fixation, and viewing of scrambled or face videos. Mean of median saccade amplitudes (left) and fixation durations (right) from 4 common marmosets following administration of saline (blue) or ketamine (right) for Fixation, Scrambled Video and Face Video blocks. \**p* < .05.

### Fixation duration

Median fixation durations were computed for each monkey (Figure 2B). A repeated measures ANOVA was conducted for the Viewing Blocks (Fixation, Scrambled Video, Face Video) and drug Treatments (Saline, Ketamine) on these data (see Figure 2b). We did not observe significant main effects of Viewing Block, *F*(2,6) = 2.04, *p* = .248, *η_p_^2^* = .405, *ε* = .502 or Treatment, *F*(1,3) = 2.64,*p* = .203, *η_p_^2^* = .468. We did observe a significant interaction of Viewing Block and Treatment, *F*(2,6) = 10.75,*p* = .045, *η_p_^2^* = .782, *ε* = .508. However, followup pairwise comparisons revealed no significant differences.

### Scan path analyses

Regions of interest (ROIs) were labelled in Face Videos as described in (see Methods>Data Analysis). The left and right eyes and the snout were defined as three regions and everywhere else was defined as the “outside” region. These ROIs were selected to mirror the eye and mouth ROIs used in human research (Souter et al., 2020; Suh et al., 2020). Saccade onset locations and landing positions while viewing these videos were then classified by ROI. We then computed multinomial logistic regression models predicting the ROI containing the landing positions of the saccade by (1) Saccade Onset Region, (2) Treatment and (3) the interaction of these terms. Additionally, we used the above ROIs to model saccades while the subjects viewed the corresponding scrambled videos. Briefly, a logistic regression can be used to model a binary dependent variable by estimating the logarithm of the odds (i.e., log-odds) of one outcome over another as a linear combination of one or more independent variables. A multinomial logistic regression extends this method to a categorically distributed dependent variable with *k* possible outcomes (e.g., landing position ROI, *k* = 4) by separately estimating the log-odds of *k* – 1 outcomes (e.g., the Snout ROI) over a selected “pivot” outcome (i.e., the Outside ROI) as a linear combination of a set of independent variables (e.g., Saccade Onset Region, Treatment and the interaction of these terms) (Agresti, 2007).

Addition of the predictor Treatment to a model that contained only the intercept significantly improved fit, χ^2^(3849) = 19.2,*p* < .001. Here, ketamine injection predicted a significantly greater number of saccades terminating in the snout, but not in the eye regions (see Table 1, Figure 3A). A model with the predictor of Saccade Onset Region further significantly improved fit, χ^2^(3843) = 313 .4, *p* < .001. Here, saccades made within regions and between the eye regions were predicted as significantly more likely (see Table 2). A model with the interaction of Treatment and Saccade Onset Region significantly improved the fit, χ^2^(3831) = 24.4, *p* = .018. This interaction shows that the effect of Saccade Onset Region was significant in the saline treatment condition (see Figure 4A) but abolished when ketamine was administered (see Table 3, Figure 4B).

**Table 1.**
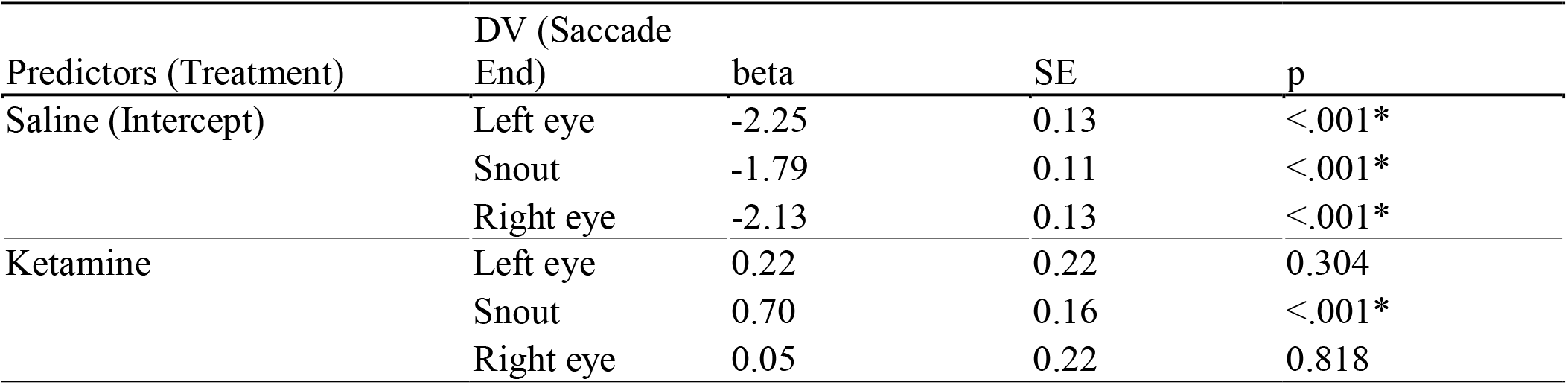
Predicting Saccade End Region with Treatment Condition using a multinomial logistic regression while marmosets viewed face videos.

**Fig 3.**
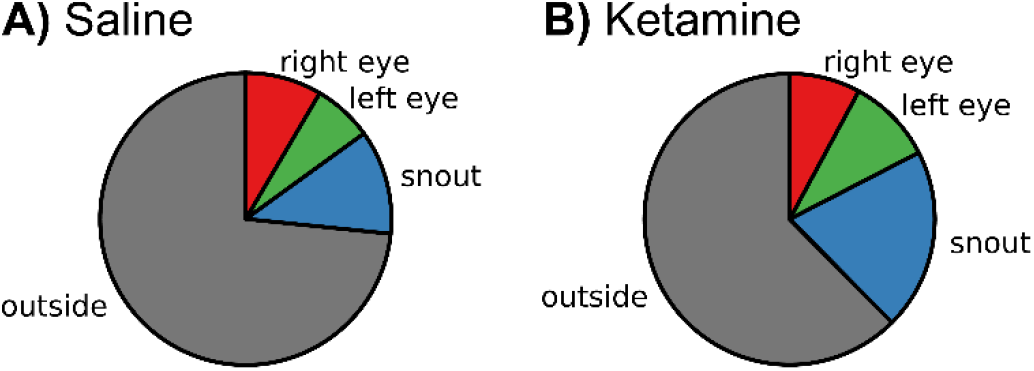
Proportions of total saccades by saccades with landing positions in each of the four regions of interest (right eye: red, left eye: green, snout: blue, outside: grey) while viewing Face Videos for saline (A) and ketamine (B) treatments.

**Table 2.** Predicting Saccade End Region with Saccade Onset Region using a multinomial logistic regression while marmosets viewed face videos.

| Predictors (Onset) | DV (Saccade End) | beta | SE | p |
| --- | --- | --- | --- | --- |
| Outside (Intercept) | Left eye | -2.48 | 0.14 | <.001* |
|  | Snout | -2.23 | 0.12 | <.001* |
|  | Right eye | -2.69 | 0.15 | <.001* |
| Left eye | Left eye | 2.00 | 0.28 | <.001* |
|  | Snout | 0.06 | 0.49 | 0.909 |
|  | Right eye | 1.30 | 0.37 | <.001* |
| Snout | Left eye | -0.29 | 0.48 | 0.547 |
|  | Snout | 2.52 | 0.19 | <.001* |
|  | Right eye | 0.51 | 0.38 | 0.186 |
| Right eye | Left eye | 0.85 | 0.37 | 0.021* |
|  | Snout | 0.24 | 0.42 | 0.561 |
|  | Right eye | 2.42 | 0.26 | <.001* |

**Fig 4.**
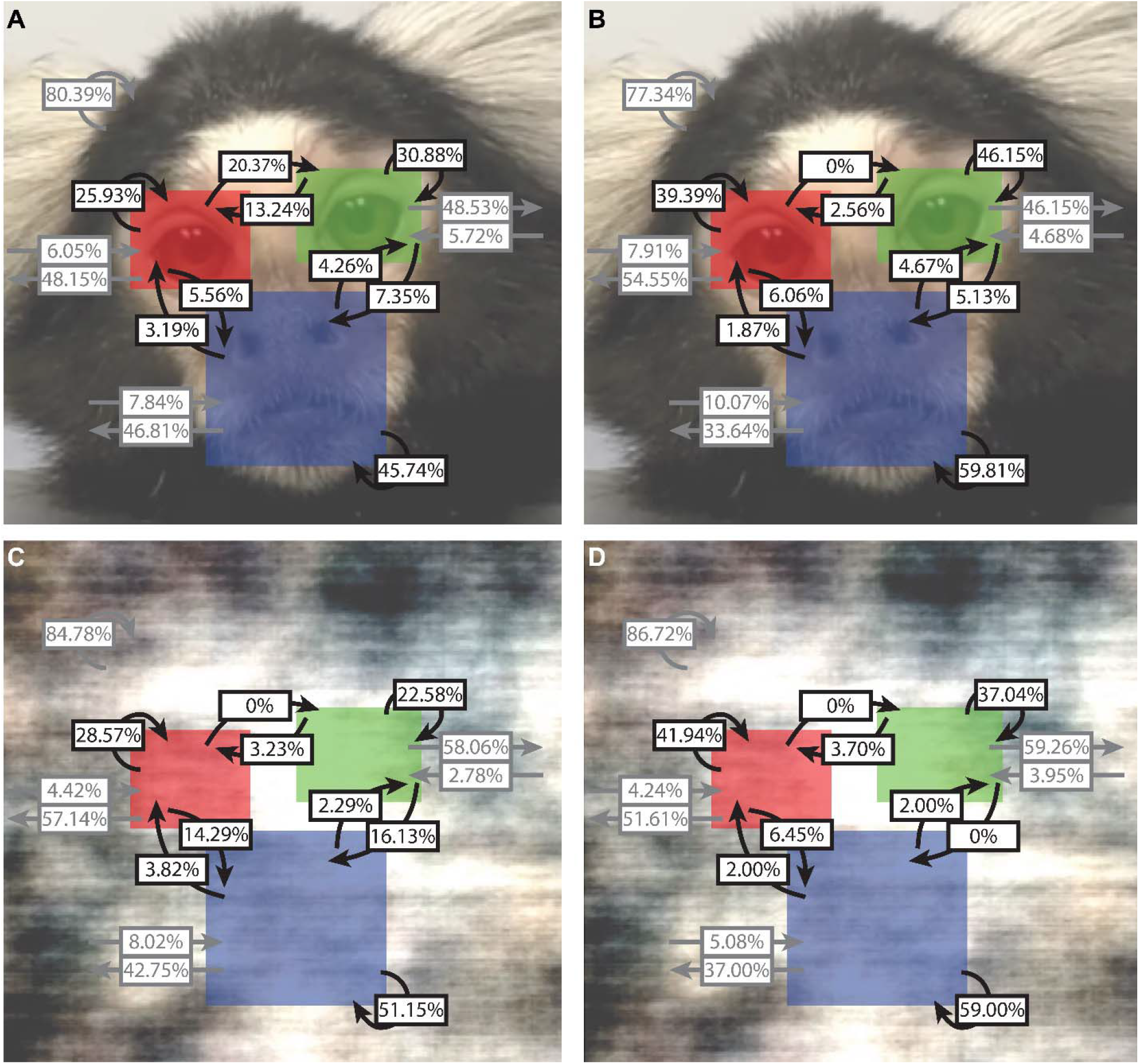
Transition probabilities of saccades between/within regions of interest during viewing of Face Videos. Proportions of saccades by region containing saccade landing positions for each saccade onset region separately for each combination of Viewing Block (unscrambled: top; scrambled: bottom) and Treatment Condition (saline: left, ketamine: right). Black arrows indicate saccades between/within facial regions (i.e., left eye: green, right eye: red, or snout: blue). Grey arrows indicate saccades to/from the outside region.

**Table 3.**
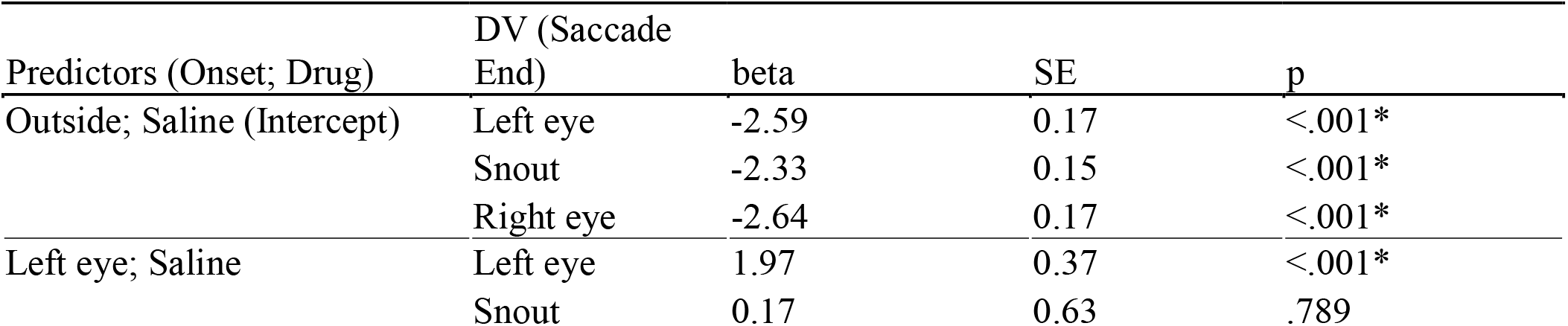
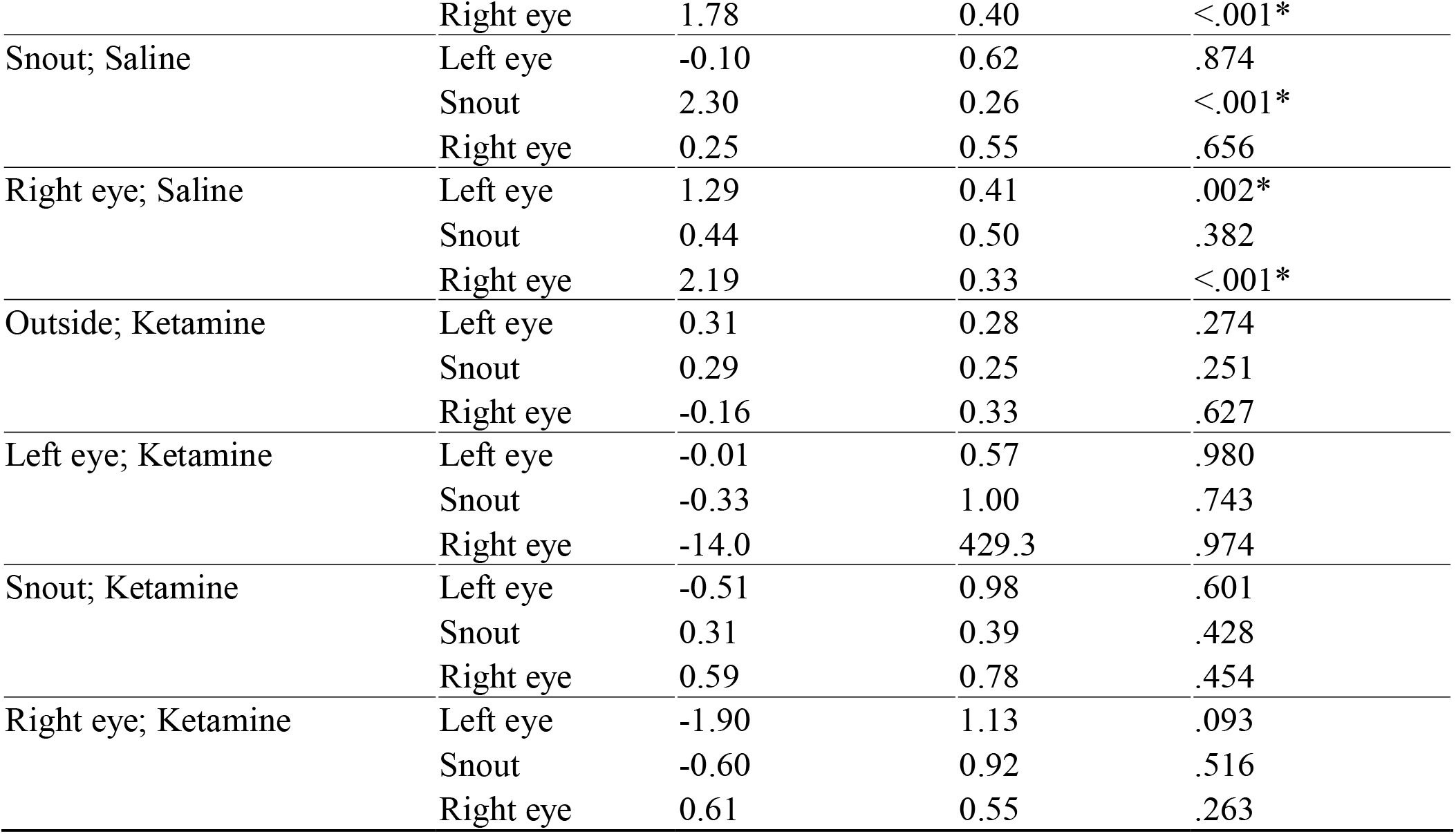
Predicting Saccade End Region with the interaction of Saccade Onset Region and Treatment Condition using a multinomial logistic regression while marmosets viewed face videos.

Conversely, when this model was constructed using saccades from the Scrambled Video Block, no significant effect of Treatment was observed, χ^2^(3996) = 2.72, *p* = .437. The model with the predictor of Saccade Onset Region significantly improved fit, χ^2^(3990) = 367.1,*p* < .001, predicting saccades within regions as more likely (see Table 4, Figure 3B). No significant interaction of Treatment and Saccade Onset Region was observed, χ^2^(3978) = 15.3, *p* = .228 (see Figure 4C,D).

**Table 4.**
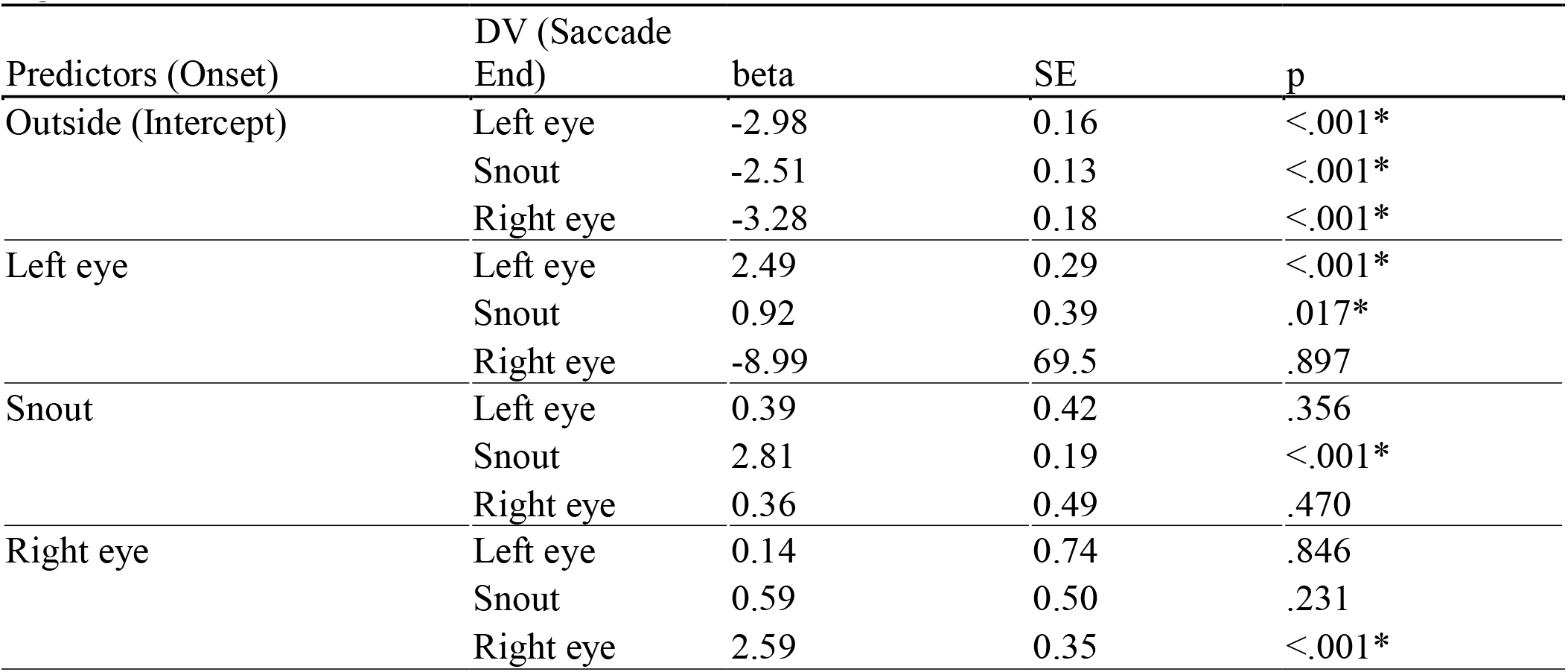
Predicting Saccade End Region with Treatment Condition using a multinomial logistic regression while marmosets viewed scrambled videos.

In sum, while the subjects viewed videos of conspecific faces, ketamine administration induced a preference for viewing the snout over the eyes. Further, saccades within a region or between the eyes were more likely. However, this structure was disrupted by ketamine administration. These effects are not observed when the subject viewed scrambled versions of these videos.

## Discussion

The marmoset model holds substantial promise for investigations of neural circuits underlying social behaviour. These highly social animals possess oculomotor behaviour and responses to face stimuli similar to those seen in macaques and humans. Recent work shows marmosets have a conserved face processing network resembling the “face patches” observed in macaques and humans (Hung et al., 2015a, 2015b; Schaeffer et al., 2020). This presents an opportunity to use these nonhuman primates for investigations of face processing and how it is disrupted in disease states. To this end, we used subanesthetic dose ketamine injections to simulate symptoms observed in neuropsychiatric disorders such as schizophrenia and observed the oculomotor behaviour of marmosets viewing videos of conspecific faces. We found that saccade amplitudes and fixation durations were not significantly altered by ketamine while viewing faces, demonstrating that the low doses of ketamine had limited effects on saccade control in general. However, significant ketamine-induced disruptions of scan paths were observed during viewing of conspecific faces, but not scrambled versions of these faces. We observed a significant difference in the distribution of saccades to the snout but not eye regions following ketamine as compared to saline. Further, saccades within regions, and between the eye regions were predicted as significantly more likely when marmosets were treated with saline, but these patterns were abolished when treated with ketamine.

Subanaesthetic doses of ketamine have been previously shown to have effects on low-level oculomotor behaviour. Ketamine elicits spontaneous nystagmus in humans (Elia & Tramèr, 2005), monkeys (Leopold et al., 2002) and cats (Godaux et al., 1990). Further, Leopold and colleagues (2002) observed spontaneous eye movements following ketamine injection, and noted an impairment in gaze holding, as well as reduced frequency, amplitude, and peak velocity of saccades. In the present work, we observed minimal nystagmus and a strong reduction of saccade amplitude when the animals were presented with the unstructured scrambled visual stimuli, but these effects were attenuated when they were presented with a central fixation stimulus or video of a conspecific’s face.

Fewer investigations have been conducted with regards to the effects of ketamine on face processing. Such effects may be classified as general impairments in configural/featural processing of faces, impairments in facial emotional processing, or abnormal patterns of gaze. Here, we discuss these effects following administration of ketamine and similar observations in individuals with schizophrenia. Neill and colleagues (2015) demonstrated the absence of the facial inversion effect in humans following ketamine injections as compared to placebo treatments. The facial inversion effect is an index of configural face processing, reflecting the increased processing time required for inverted as compared to upright faces, for which strong expectations exist regarding configural information (Yin, 1969). Similar impairments of configural processing, but not featural processing, have been observed in patients with schizophrenia. While these impairments can be attributed in part to cognitive impairments and general deficits in early stages of visual processing in these patients (Butler et al., 2008; Soria Bauser et al., 2012), an additional face-specific deficit is observed (Chen et al., 2009) (c.f. Bortolon et al., 2015). In addition to general impairments in face processing, ketamine-induced impairments in facial emotion processing have been demonstrated using behavioural (Malhotra et al., 1997) and electrophysiological (Lundin et al., 2020) approaches. Such impairments have also been observed in patients with schizophrenia (Chan et al., 2010). A general difficulty with identifying the emotion being expressed has long been observed (see for review: Morris et al., 2009). In addition, a specific impairment in identifying negative emotions, such as sadness or fear, has been observed (Edwards et al., 2001; Hall et al., 2008). Neural correlates of this can be observed in the amygdala, where seemingly low activation for negative faces can be seen due to abnormally high activation for neutral faces (Hall et al., 2008; Holt et al., 2006; see for review: Aleman & Kahn, 2005). The amygdala has an established role in fear processing and is known to modulate activity in the fusiform gyrus in relation to emotional information (Rotshtein et al., 2001). To our knowledge, no other studies have been conducted investigating the effects of ketamine on scan paths of animals looking at conspecifics’ faces. However, abnormal gaze patterns, including avoidance of the eyes, is observed in patients with schizophrenia (Loughland et al., 2002; Phillips & David, 1998; Williams et al., 1999); damage to the amygdala has been documented reducing eye contact in human’s engaging in conversations with real people (Spezio et al., 2007). Scan path abnormalities are also observed in individuals with ASD, though the effect has been shown to be restricted to reduction in eye contact (Papagiannopoulou et al., 2014; Yi et al., 2013).

The eyes are an important facial feature and it has been demonstrated that fixations on eyes are predictive of increased face identification accuracy and a reduction of the facial inversion effect (Hills et al., 2013). Intranasal administration of oxytocin, a hormone with a role in enhancing prosocial behaviours, has been shown to increase fixations to eye regions relative to the mouth region in macaques (Dal Monte et al., 2014) and marmosets (Kotani et al., 2017) demonstrating the value of this metric in assessing primate scanning behaviour of conspecific faces (see also: Souter et al., 2020). In the present work, we observed a pattern consistent with observations in patients with neuropsychiatric disorders; probability of saccades increased to the snout region but not to eye regions following injection with ketamine, effectively reducing the ratio of time spent on eyes vs the snout. It should be noted that fewer saccades were made to non-face regions following ketamine injection, though this is likely attributable to the fact that faces were presented at the center of the screen and fewer large saccades were made by the subjects following ketamine administration. However, this central fixation bias alone cannot fully explain our observations as this effect was not present for scrambled versions of the videos.

Recent work has also investigated visual exploration of faces in humans by investigating the probabilities of transitions between regions of interest and observed that individuals with schizophrenia were less likely to transition from the mouth to the eye region (Suh et al., 2020). In the present work, when monkeys were injected with saline, we observed a gaze pattern in which saccades within a region on the face and between the eyes are more probable. However, this pattern was lost following ketamine administration. Investigations in patients have revealed that these abnormal gaze patterns are most pronounced in free viewing tasks as we used here, and more closely resemble healthy controls when in a task, such as identifying the age or gender of the imaged individual (Delerue et al., 2010). Future investigations using marmosets trained on a formal task, such as a match-to-sample, in which second order features are manipulated, may prove illuminating in disentangling the role of ketamine on holistic face processing and provide a model to investigate the interaction of face processing and scan path abnormalities observed in neuropsychiatric disorders.

The specific contributions of certain brain areas to normal face scanning behaviour remains an area of active interest. Recent work in macaques has demonstrated a role of the orbitofrontal cortex (OFC) in scanning of faces (Goursaud & Bachevalier, 2020). Specifically, lesions in the OFC resulted increased overall looking at faces and increased attention to the eyes, which the authors posit may be due to the absence of top-down influences from OFC on sensory association cortices and subcortical structures like the amygdala, which play a role in allocating attention to social and non-social stimuli. Future pharmacological investigations targeting established nodes of the face processing network would prove invaluable in further advancing our knowledge on this topic.

In conclusion, our findings demonstrate that ketamine induces a substantial impairment of holistic face processing in the common marmoset monkey and support its use as a model investigating face processing networks and their impairments in neuropsychiatric disorders.

## Acknowledgements

We thank C. Vander Tuin, W. Froese, and K. Faubert for expert technical and surgical assistance, and care of the marmosets.

## Grants

This research was supported by the Canadian Institutes of Health Research grant FRN148365 to S.E. and the Canada First Research Excellence Fund to BrainsCAN. JS was supported by the Natural Sciences and Engineering Research Council’s Canadian Graduate Scholarship (Doctoral).

## Notes

### Competing Interest Statement

The authors have declared no competing interest.

